## Supplemental Figures and Tables for "Genomic Resources for the Scuttle Fly *Megaselia abdita*: A Model Organism for Comparative Developmental Studies in Flies"

Figures S1 – S11
Tables S1 – S4

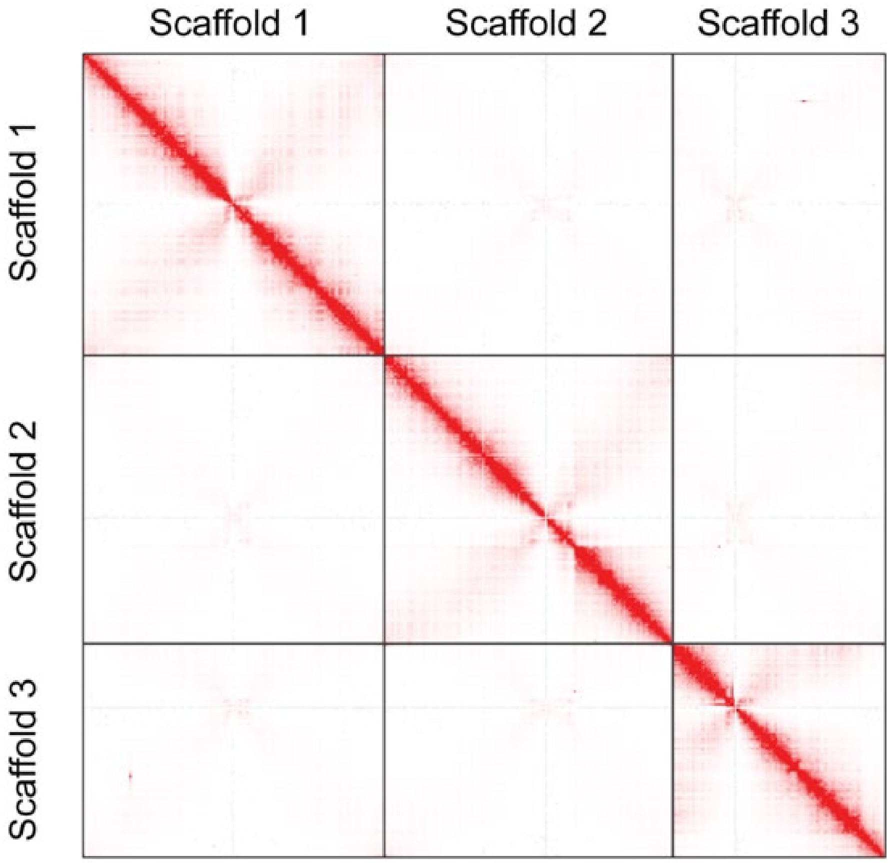


**Figure S1.** Hi-C contact map showing spatial proximity between scaffolds in the genome assembly. The intensity of red indicates the frequency of interactions between genomic regions, with darker red representing higher contact frequencies. Scaffolds are arranged along the axes, and strong diagonal signals indicate intra-scaffold interactions.

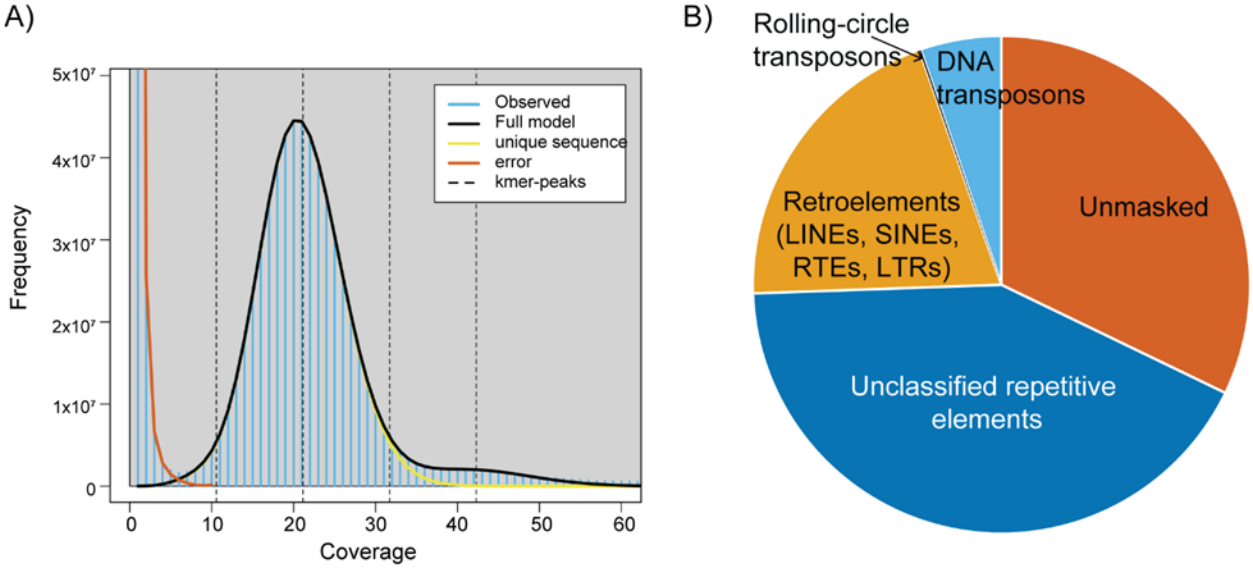


**Figure S2.** *M. abdita* genome characteristics: k-mer distribution and repetitive element composition.

A) 21-mer frequency distribution for *M. abdita* PacBio reads. The observed k-mer coverage (blue) is modeled by GenomeScope (black line), which includes contributions from unique sequences (yellow) and sequencing errors (orange).

B) Proportion of the *M. abdita* genome corresponding to repetitive elements. Retrotransposons (e.g., LINEs, SINEs, RTEs, LTRs), DNA transposons, rolling-circle transposons, unclassified repetitive elements, and unmasked regions are also included.

**Table S1**. Scaffold lengths of *M. abdita's* genome assembly.

| Scaffold | Length (bp) |
|---|---|
| 1 | 222,469,999 |
| 2 | 212,802,314 |
| 3 | 155,577,902 |
| 4 | 1,302,958 |
| 5 | 455,293 |
| 6 | 55,021 |
| 7 | 35,625 |
| 8 | 29,721 |
| 9 | 17,979 |
| 10 | 17,687 |
| 11 | 17,313 |
| 12 | 13,866 |
| 13 | 13,720 |
| 14 | 10,567 |
| 15 | 4,999 |

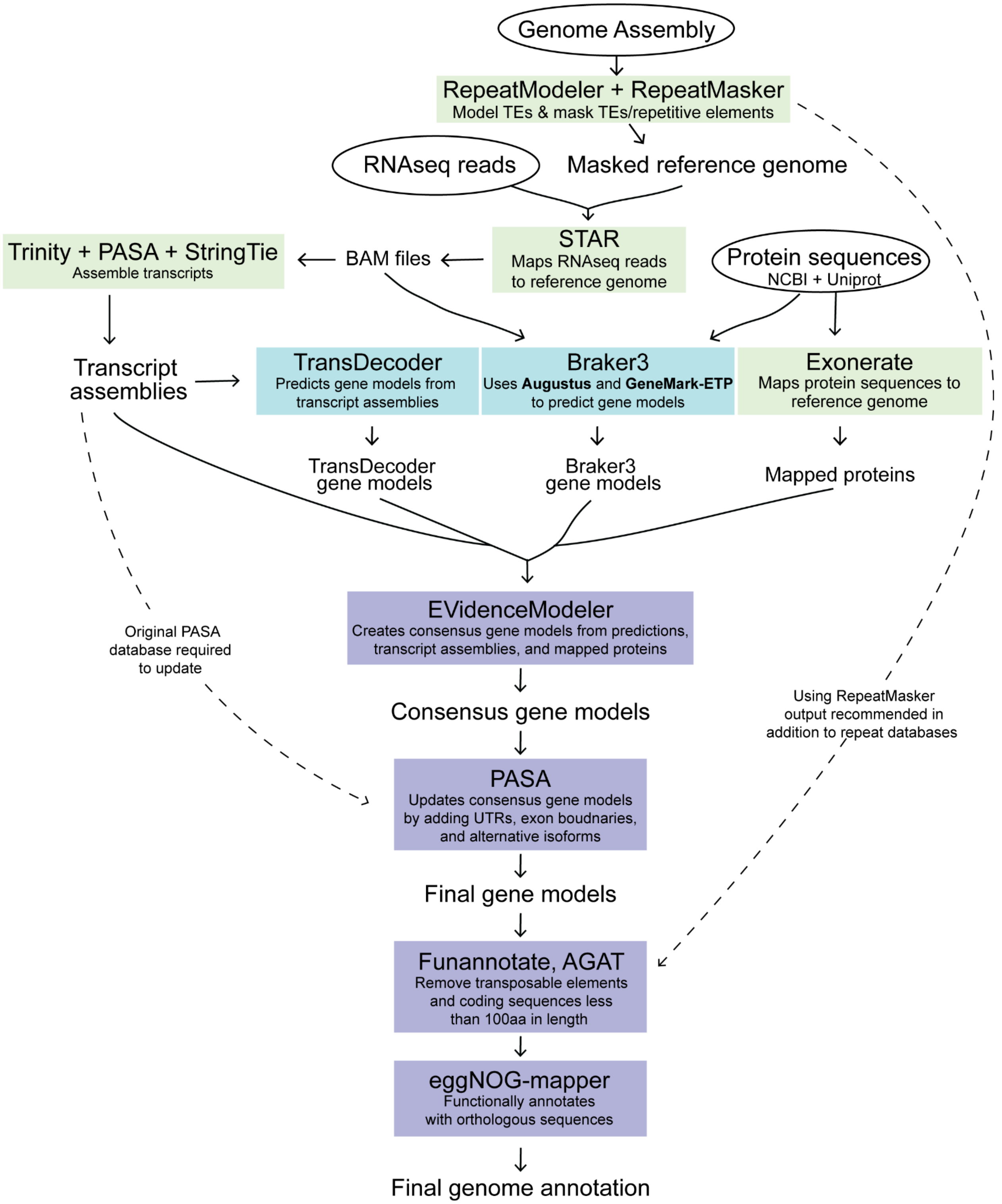


**Figure S3.** Genome annotation pipeline. The starting input files are shown in black ovals and include the genome assembly along with two lines of evidence: RNAseq reads (fastq format) and protein sequences obtained from NCBI and UniProt. Software tools are represented in colored boxes: green indicates mapping software, blue indicates gene model generation software, and purple indicates post-gene model processing and functional annotation tools.

Arrows pointing from a software box to unboxed text represent the output files generated by the software. Arrows leading from unboxed text to a software box indicate input files used by the software.

**Table S2.** Comparison of *D. melanogaster* and *M. abdita's* protein-coding genes exon, transcript, and full gene length in base pairs (bp). All *D. melanogaster* non-protein coding genes were removed before analysis.

| | Min | Median | Mean | Max |
|---|---|---|---|---|
| **Exons** | | | | |
| *D. melanogaster* | 1 | 250 | 553 | 28,074 |
| *M. abdita* | 3 | 293 | 505 | 19,709 |
| **Transcript (all isoforms)** | | | | |
| *D. melanogaster* | 152 | 1,700 | 2,353 | 71,382 |
| *M. abdita* | 306 | 1,581 | 2,053 | 43,640 |
| **Gene** | | | | |
| *D. melanogaster* | 162 | 2,159 | 6,960 | 1,965,856 |
| *M. abdita* | 306 | 6,650 | 19,479 | 476,305 |

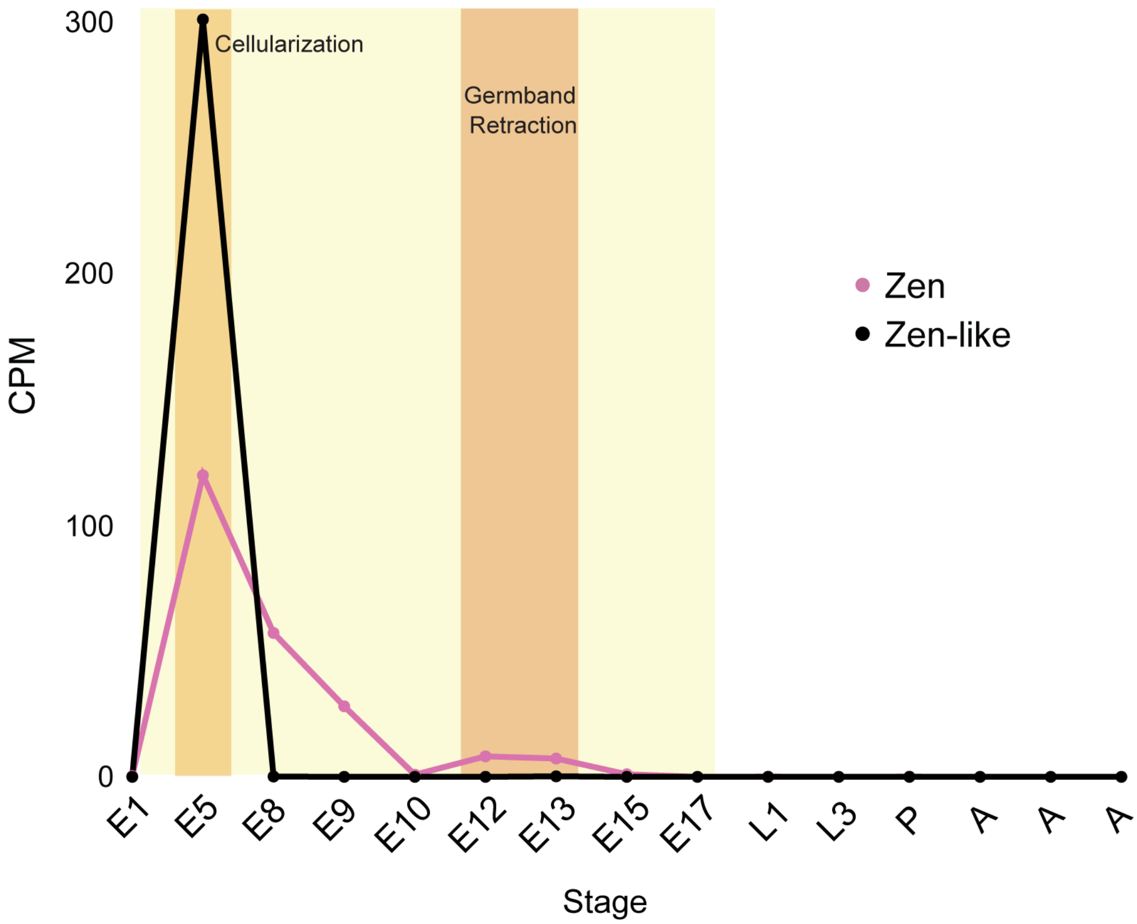


**Figure S4.** Expression of *Zen* (pink) and *Zen-like* (black) genes in *M. abdita* across developmental stages. Expression is in counts per million (CPM). Yellow shading (E1–E17) indicates embryonic stages, while L, P, and A represent larval instars, an early pupal stage, and three different adult samples, respectively. Key embryonic developmental events, cellularization and germband retraction, are highlighted.

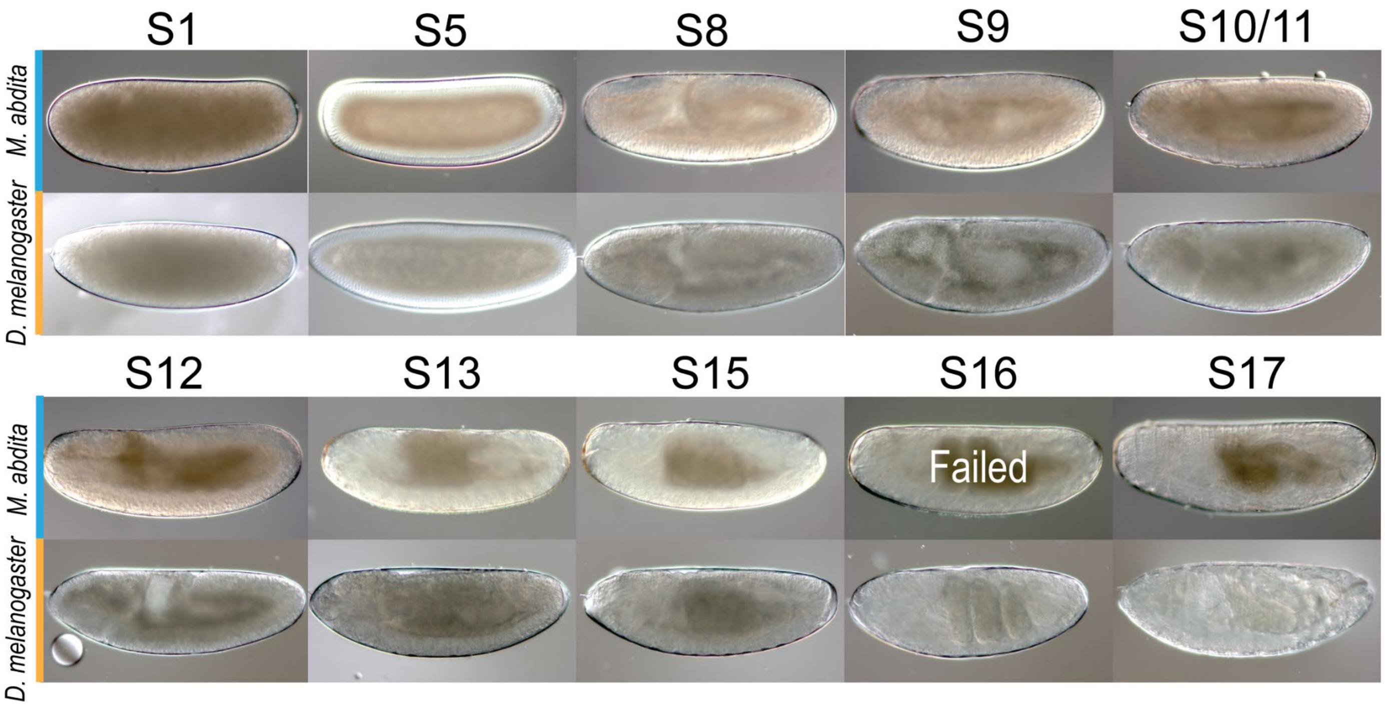


**Figure S5.** Images of the embryos used for individual RNA sequencing, spanning developmental stages 1 to 17. *M. abdita* embryos are shown in the top row (blue bar) and *D. melanogaster* on the bottom row (yellow bar). Each stage is indicated above the corresponding images. Note that for stage 16 of *M. abdita*, sequencing failed.

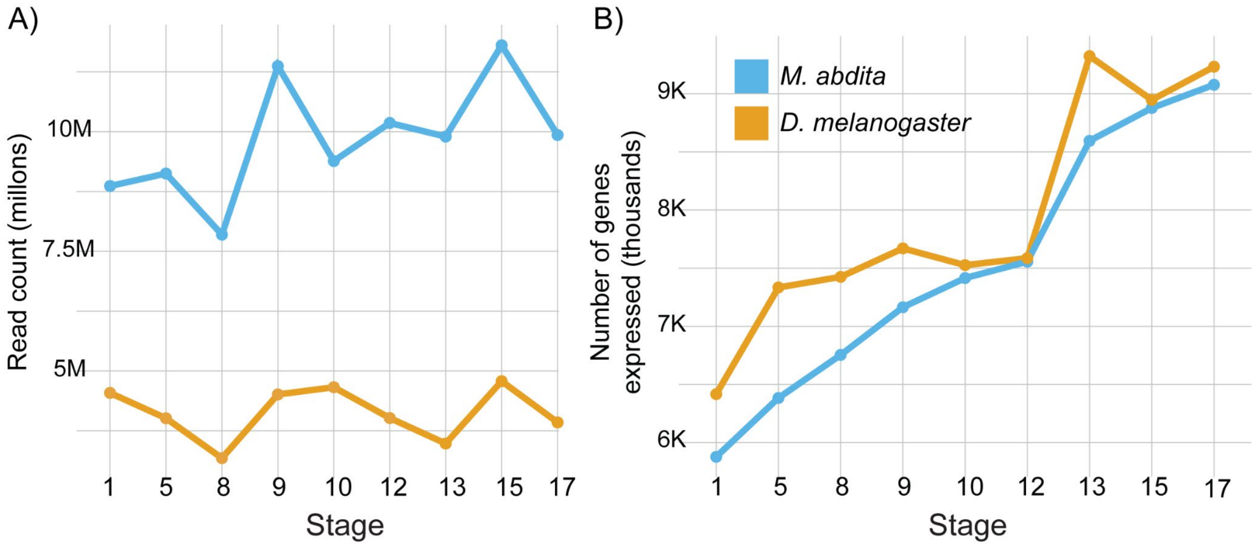


**Figure S6.** RNA-seq read counts and gene expression across embryonic stages in *D. melanogaster* and *M. abdita*

A) Comparison of raw RNA-seq read counts for single embryos of *D. melanogaster* (yellow) and *M. abdita* (blue) across embryonic developmental stages.

B) Comparison of the number of genes expressed in single embryos of *D. melanogaster* (yellow) and *M. abdita* (blue) across embryonic developmental stages.

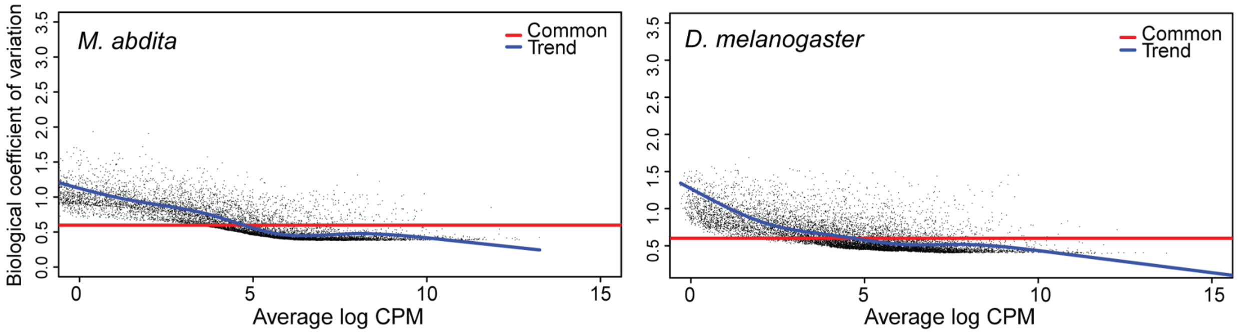


**Figure S7.** Biological coefficient of variation (BCV) plots for *M. abdita* (left) and *D. melanogaster* (right) RNA-seq data. Each black dot represents a gene, with the BCV plotted against the average log counts per million ($log_2$CPM). The red line indicates the global common dispersion ($BCV^2$ is equal to dispersion), while the blue line shows the trend of gene-specific dispersions as a function of expression level.

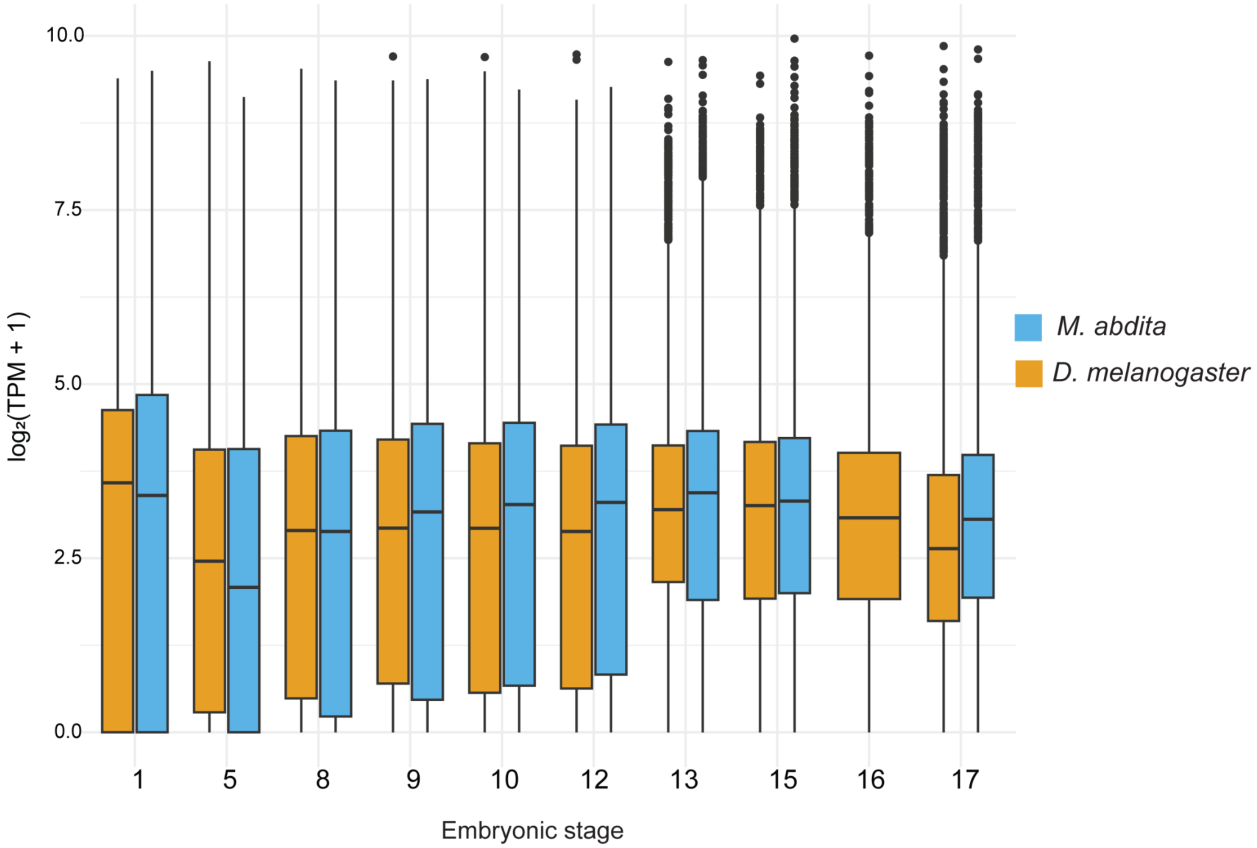


**Figure S8**. Boxplot of gene expression in *D. melanogaster* (yellow) and *M. abdita* (blue) by embryonic stage. The y-axis represents log-transformed transcript abundance (TPM), and the x-axis represents the developmental stages. The boxes show the interquartile range (IQR), with the median indicated by the horizontal line. Black dots indicate outliers.

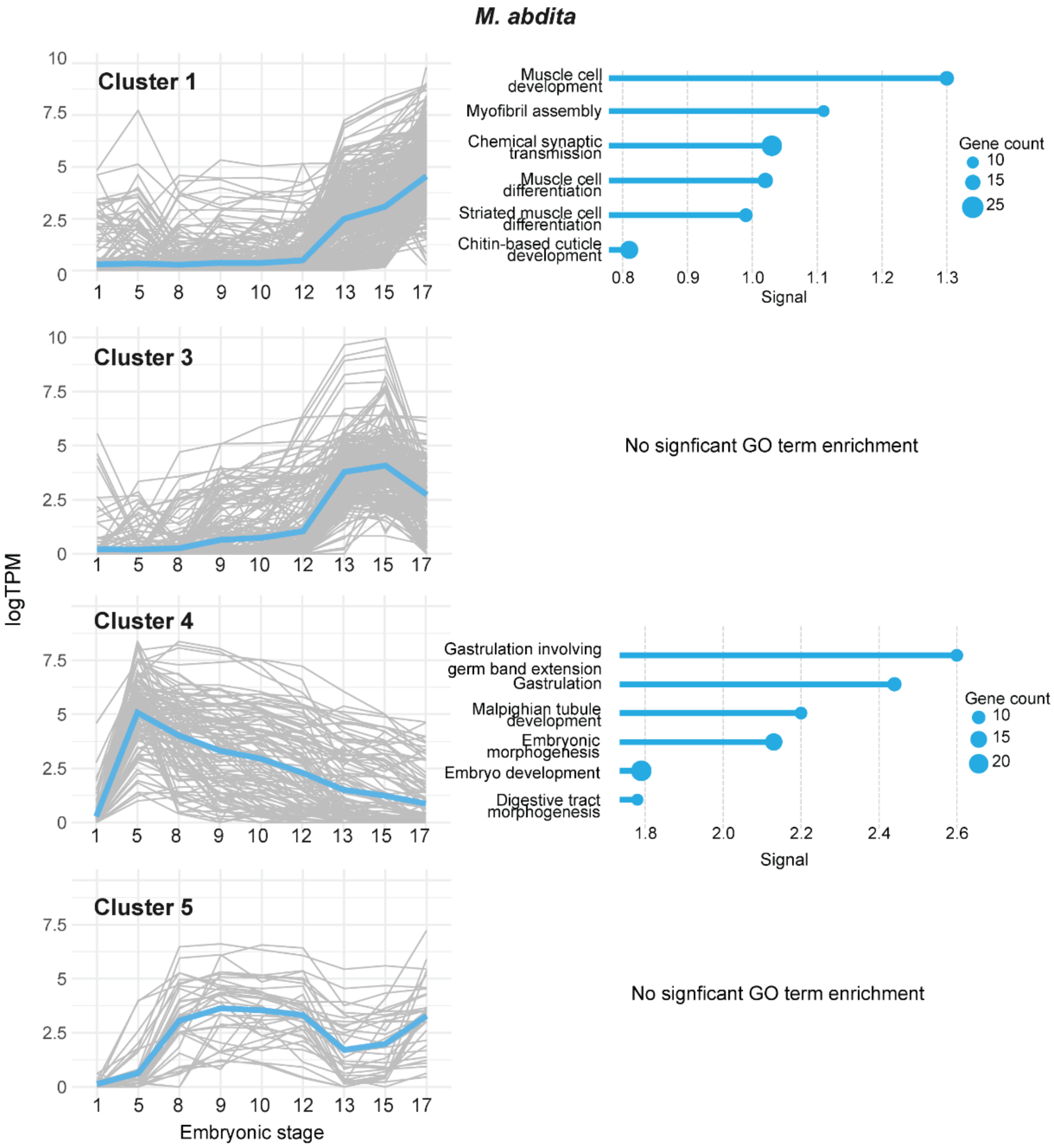
M. abdita
Cluster 1
Cluster 3
Cluster 4
Cluster 5
logTPM
Embryonic stage
Muscle cell development
Myofibril assembly
Chemical synaptic transmission
Muscle cell differentiation
Striated muscle cell differentiation
Chitin-based cuticle development
Signal
Gene count
No signficant GO term enrichment
Gastrulation involving germ band extension
Gastrulation
Malpighian tubule development
Embryonic morphogenesis
Embryo development
Digestive tract morphogenesis
Signal
Gene count
No signficant GO term enrichment

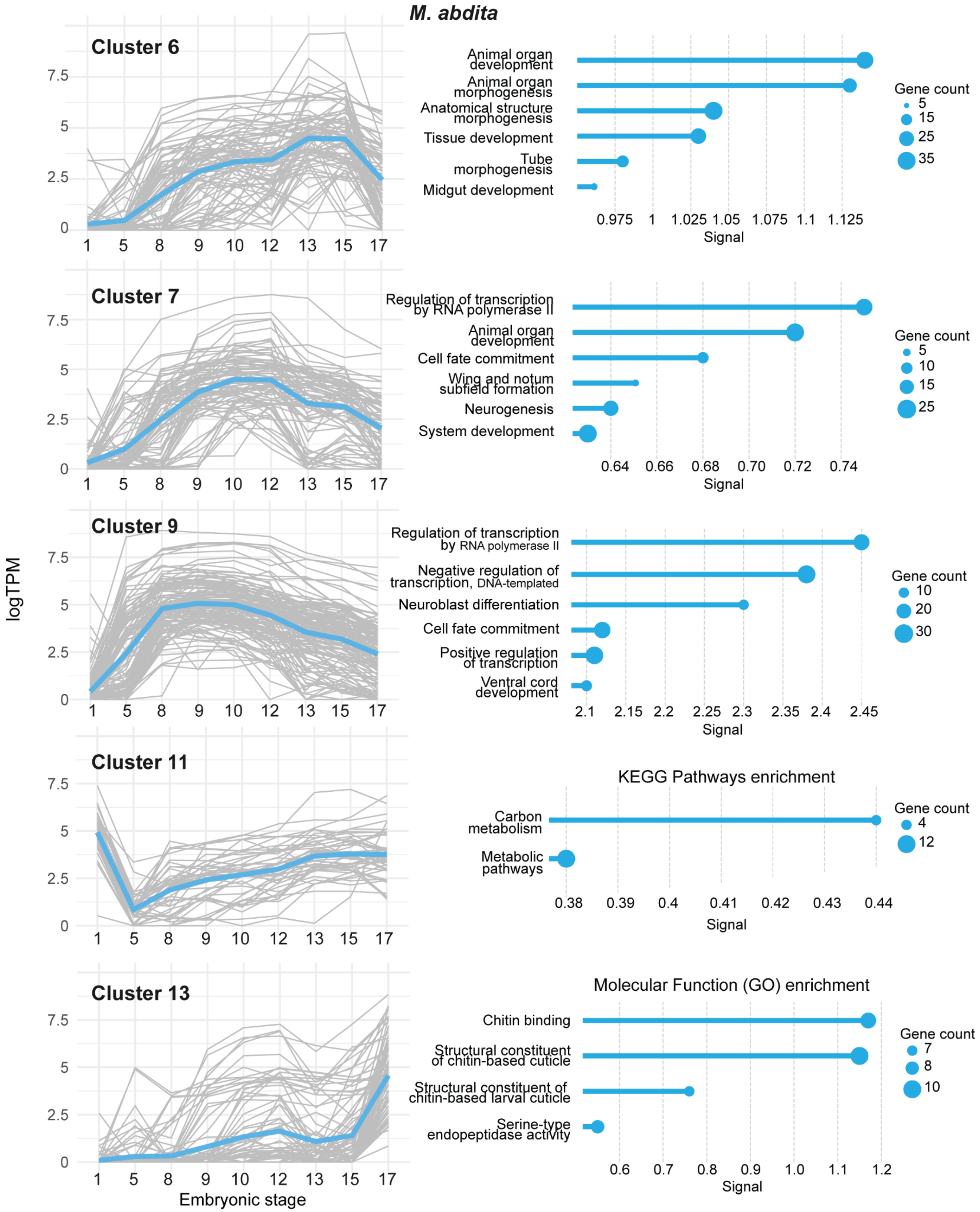
M. abdita
Cluster 6
Cluster 7
Cluster 9
Cluster 11
Cluster 13
logTPM
Embryonic stage
Animal organ development
Animal organ morphogenesis
Anatomical structure morphogenesis
Tissue development
Tube morphogenesis
Midgut development
Regulation of transcription by RNA polymerase II
Cell fate commitment
Wing and notum subfield formation
Neurogenesis
System development
Negative regulation of transcription, DNA-templated
Neuroblast differentiation
Positive regulation of transcription
Ventral cord development
KEGG Pathways enrichment
Carbon metabolism
Metabolic pathways
Molecular Function (GO) enrichment
Chitin binding
Structural constituent of chitin-based cuticle
Structural constituent of chitin-based larval cuticle
Serine-type endopeptidase activity
Gene count
Signal

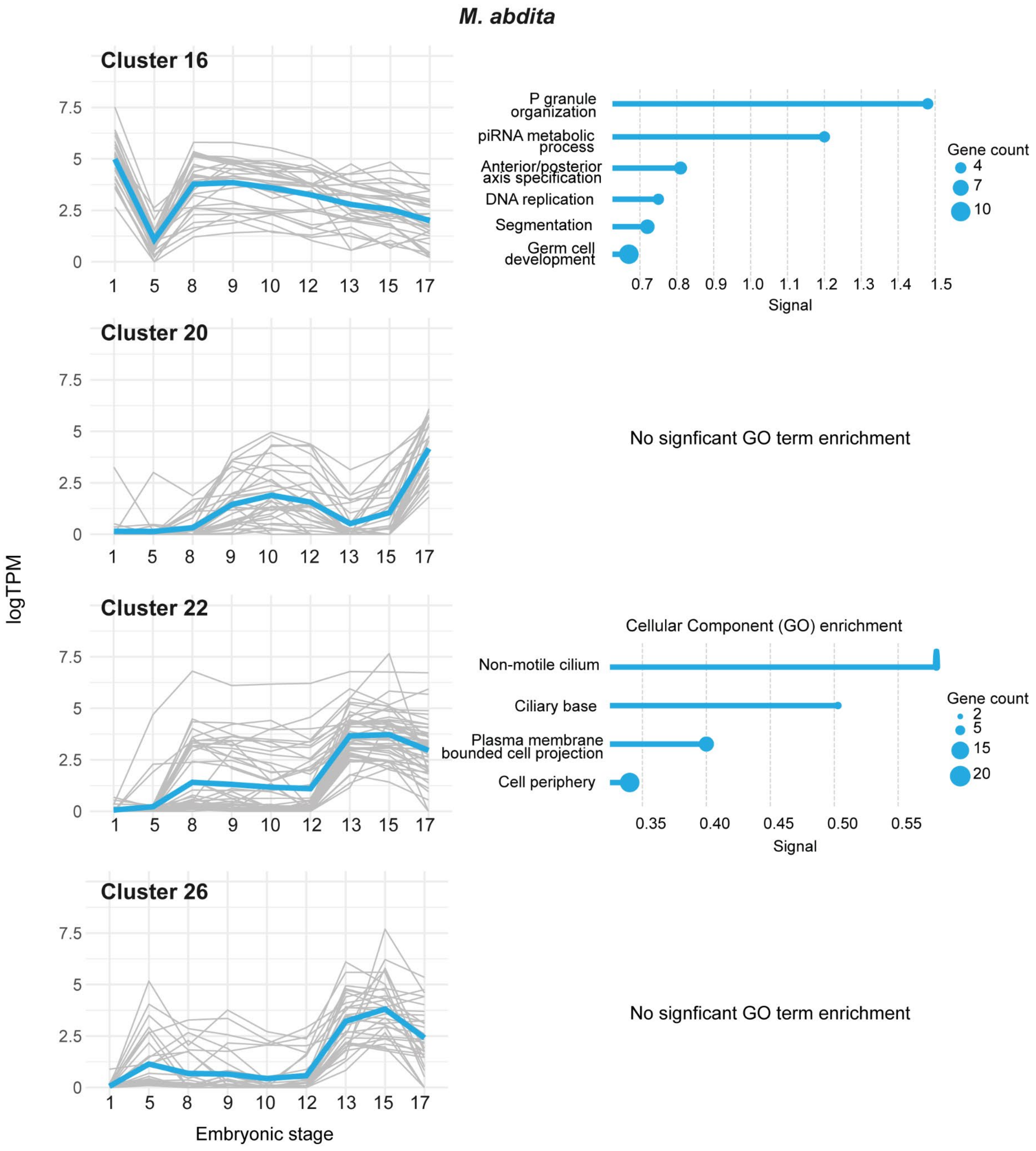
M. abdita
Cluster 16
Cluster 20
Cluster 22
Cluster 26
logTPM
Embryonic stage
P granule organization
piRNA metabolic process
Anterior/posterior axis specification
DNA replication
Segmentation
Germ cell development
Signal
Gene count
No signficant GO term enrichment
Cellular Component (GO) enrichment
Non-motile cilium
Ciliary base
Plasma membrane bounded cell projection
Cell periphery
Signal
Gene count
No signficant GO term enrichment

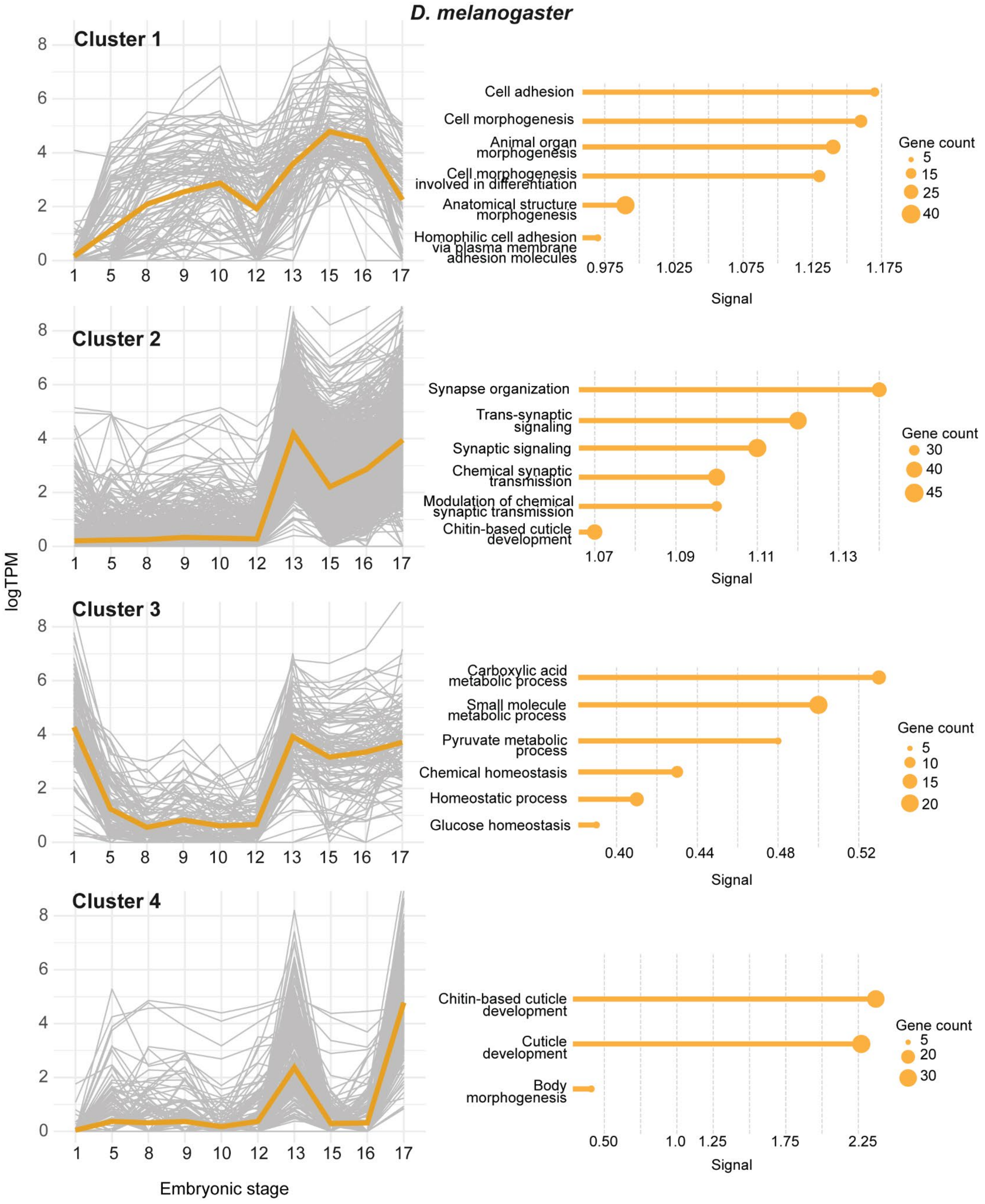

D. melanogaster
Cluster 1
Cluster 2
Cluster 3
Cluster 4
logTPM
Embryonic stage
Cell adhesion
Cell morphogenesis
Animal organ morphogenesis
Cell morphogenesis involved in differentiation
Anatomical structure morphogenesis
Homophilic cell adhesion via plasma membrane adhesion molecules
Synapse organization
Trans-synaptic signaling
Synaptic signaling
Chemical synaptic transmission
Modulation of chemical synaptic transmission
Chitin-based cuticle development
Carboxylic acid metabolic process
Small molecule metabolic process
Pyruvate metabolic process
Chemical homeostasis
Homeostatic process
Glucose homeostasis
Chitin-based cuticle development
Cuticle development
Body morphogenesis
Signal
Gene count

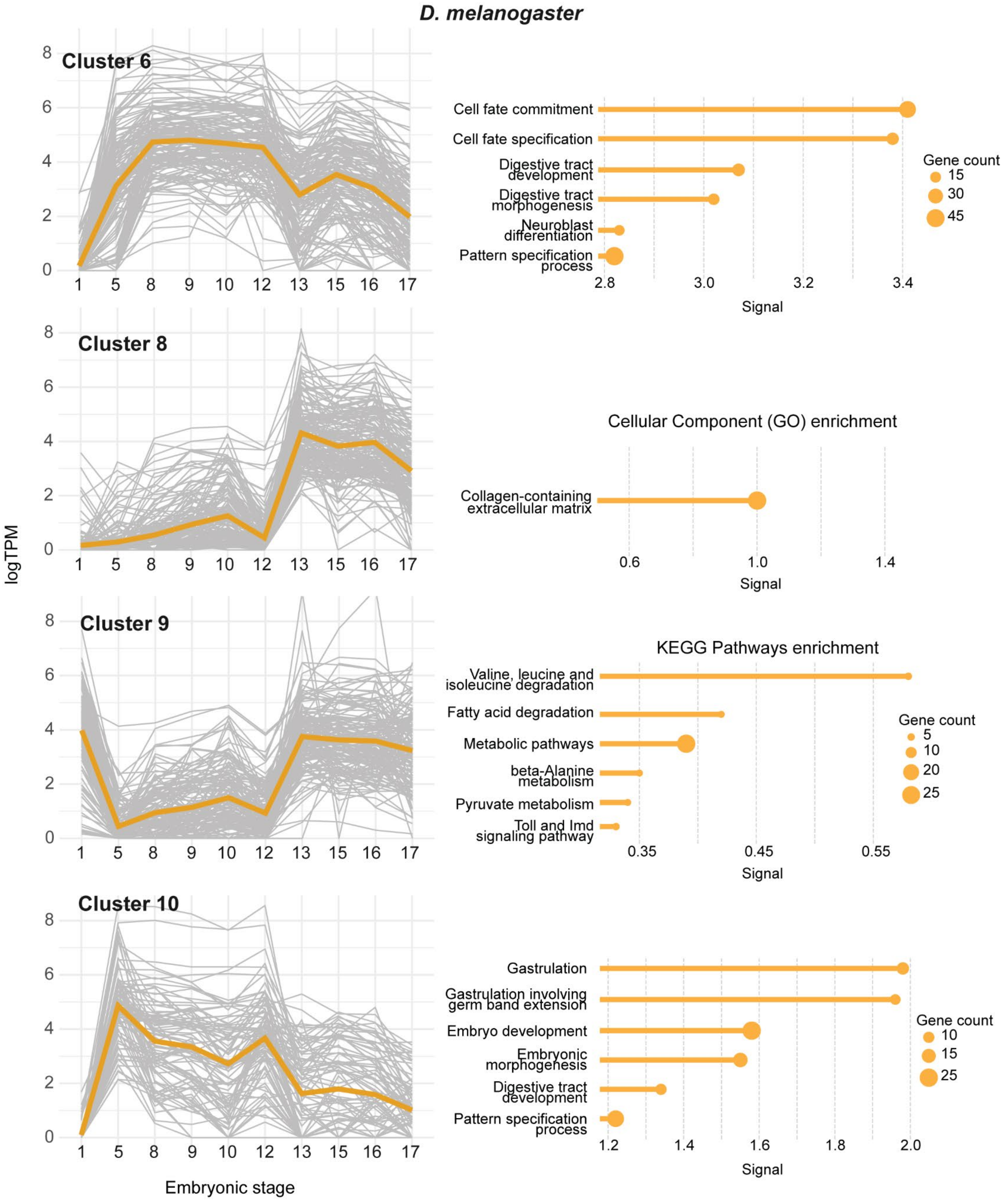

D. melanogaster
Cluster 6
Cluster 8
Cluster 9
Cluster 10
logTPM
Embryonic stage
Cell fate commitment
Cell fate specification
Digestive tract development
Digestive tract morphogenesis
Neuroblast differentiation
Pattern specification process
Gene count
15
30
45
Signal
Cellular Component (GO) enrichment
Collagen-containing extracellular matrix
KEGG Pathways enrichment
Valine, leucine and isoleucine degradation
Fatty acid degradation
Metabolic pathways
beta-Alanine metabolism
Pyruvate metabolism
Toll and Imd signaling pathway
Gene count
5
10
20
25
Gastrulation
Gastrulation involving germ band extension
Embryo development
Embryonic morphogenesis
Digestive tract development
Pattern specification process
Gene count
10
15
25

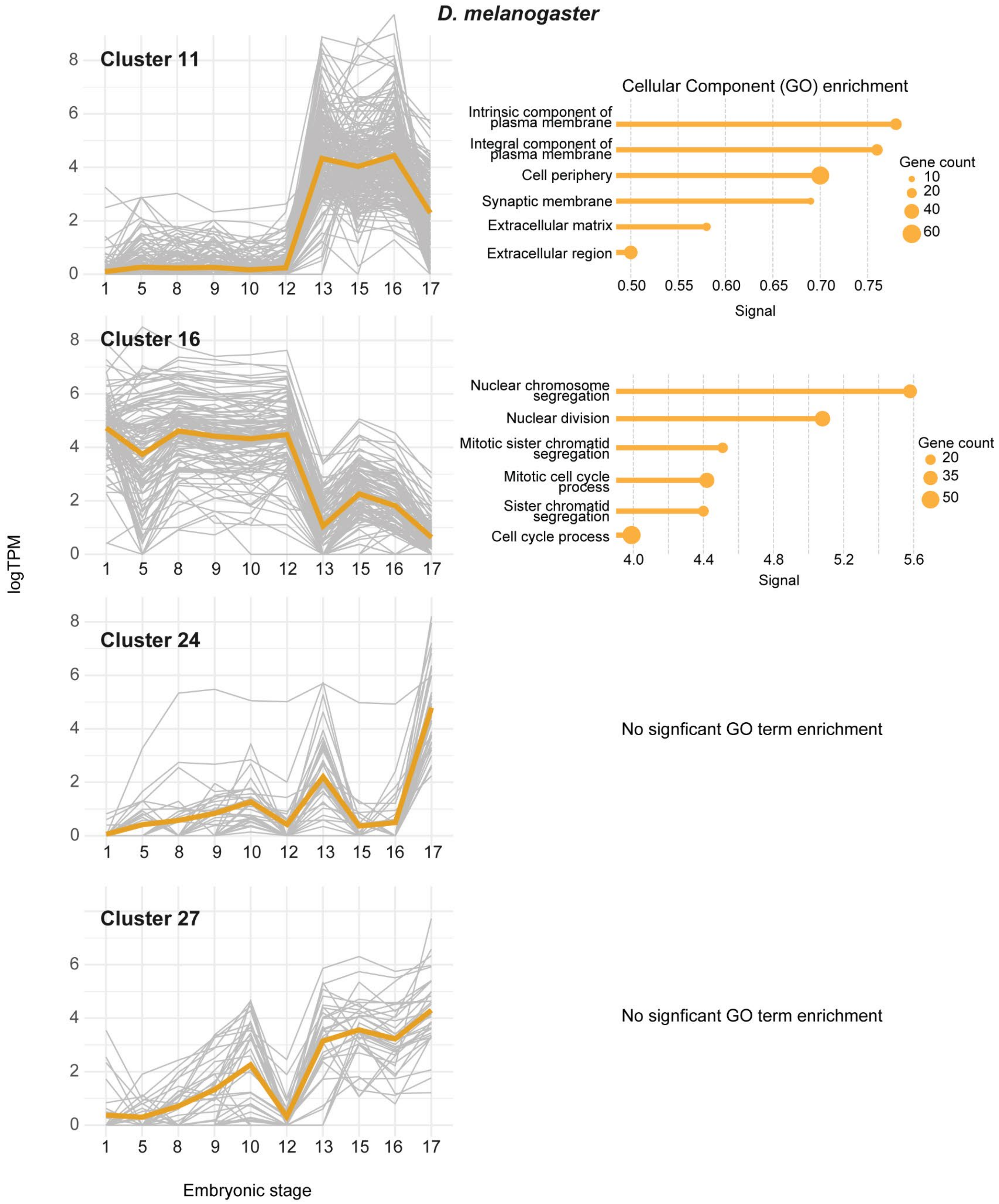

D. melanogaster
Cluster 11
Cellular Component (GO) enrichment
Intrinsic component of plasma membrane
Integral component of plasma membrane
Cell periphery
Synaptic membrane
Extracellular matrix
Extracellular region
Gene count
10
20
40
60
0.50
0.55
0.60
0.65
0.70
0.75
Signal
Cluster 16
Nuclear chromosome segregation
Nuclear division
Mitotic sister chromatid segregation
Mitotic cell cycle process
Sister chromatid segregation
Cell cycle process
Gene count
20
35
50
4.0
4.4
4.8
5.2
5.6
Signal
Cluster 24
No signficant GO term enrichment
Cluster 27
No signficant GO term enrichment
logTPM
1 5 8 9 10 12 13 15 16 17
Embryonic stage

**Figure S9**. Differentially expressed gene clusters and enriched functional annotations during embryogenesis in *M. abdita* and *D. melanogaster*.

(Left) Plots of differentially expressed gene (DEG) clusters during embryogenesis. *M. abdita* clusters are shown in blue and *D. melanogaster* clusters in yellow. Grey lines indicate expression across embryonic stages for individual genes, with the average expression profile shown in bold for each group of genes. DEGreports generated cluster names (e.g., "Cluster 1") which are arbitrary but retained for continuity.

(Right) Each cluster's enriched Biological Process Gene Ontology (BP GO) terms; if there were no enriched BP GO terms, enriched Cellular component (CC) or KEGG pathway terms are shown. All GO terms are ranked by signal which is defined as the weighted harmonic mean between the observed/expected ratio of genes belonging to a specific GO term and -log(FDR). Since lower FDRs bias towards larger GO terms and the observed/expected ratio toward smaller GO terms, signal attempts to balance both metrics in ranking GO terms.

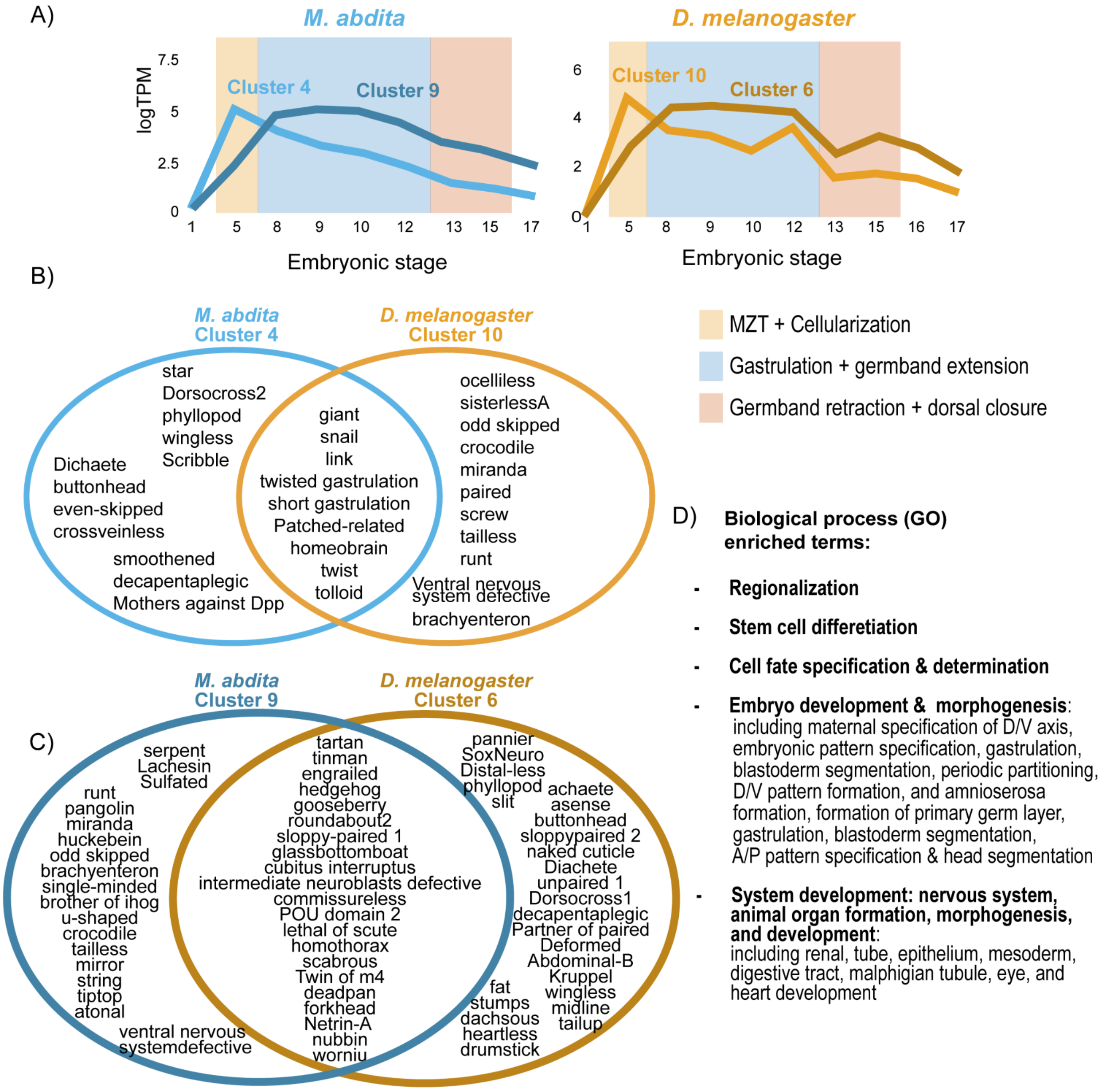


**Figure S10.** Differentially expressed gene clusters in *M. abdita* and *D. melanogaster* enriched for development gene ontology terms. A) Average expression profiles (log2(TPM + 1)) for genes in *M. abdita* clusters 4 and 9 (left) and *D. melanogaster* clusters 10 and 6 (right). Developmental stages are indicated on the x-axis, with key embryonic events (MZT + cellularization, gastrulation + germband extension, and germband retraction + dorsal closure) highlighted. B) Venn diagram showing named differentially expressed genes (DEGs) shared between *M. abdita* cluster 4 and *D. melanogaster* cluster 10.

C) Venn diagram showing named DEGs shared between *M. abdita* cluster 9 and *D. melanogaster* cluster 6.

D) Shared enriched Biological Process Gene Ontology (GO) terms for all clusters.

**Table S3.** Coding and protein sequence characteristics of orphan genes in *M. abdita*. The table includes coding sequence (nucleotide) and amino acid sequence (protein) characteristics for each orphan gene (Gene ID as it appears in the genome annotation). Genes highlighted in blue have an amino acid sequence with a stability index classifying them as "stable." "n.s." indicates "no significant match" in sequence similarity searches using NCBI's blast.

| | Nucleotide | | | | | Protein | | | | |
|---|---|---|---|---|---|---|---|---|---|---|
| Gene ID | ORF length (AA) | Fickett Score | Isoelectric Point | Coding Probability | nblast | alphafold (pTM) | Instability Index | Aliphatic Index | Average of hydropathicity | pblast |
| evm.TU.Scaffold_3_155577901.980 | 571 | 0.41 | 6.46 | 1.00 | n.s. | 0.57 | 38.92 | 99.51 | -0.12 | n.s. |
| evm.TU.Scaffold_1_222469999.1571 | 174 | 0.39 | 6.09 | 0.98 | n.s. | 0.53 | 36.79 | 85.66 | -0.44 | n.s. |
| evm.TU.Scaffold_1_222469999.2459 | 102 | 0.46 | 8.08 | 0.53 | n.s. | 0.39 | 30.12 | 115.74 | 0.60 | n.s. |
| evm.TU.Scaffold_2_212802314.799 | 139 | 0.45 | 5.06 | 0.97 | n.s. | 0.36 | 37.03 | 77.75 | -0.03 | n.s. |
| evm.TU.Scaffold_1_222469999.1775 | 105 | 0.44 | 4.54 | 0.79 | n.s. | 0.32 | 28.32 | 139.71 | 0.82 | n.s. |
| evm.TU.Scaffold_1_222469999.6 | 264 | 0.43 | 9.66 | 1.00 | n.s. | 0.29 | 29.60 | 76.84 | -0.15 | n.s. |
| evm.TU.Scaffold_2_212802314.3488 | 181 | 0.44 | 4.32 | 1.00 | n.s. | 0.21 | 19.35 | 88.89 | -0.04 | n.s. |
| evm.TU.Scaffold_3_55577901.555 | 459 | 0.36 | 5.30 | 1.00 | n.s. | 0.19 | 39.71 | 64.02 | -0.73 | n.s. |
| evm.TU.Scaffold_1_22469999.3564 | 270 | 0.43 | 9.08 | 1.00 | n.s. | 0.16 | 36.66 | 82.90 | -0.23 | n.s. |
| evm.TU.Scaffold_1_222469999.1521 | 281 | 0.42 | 9.57 | 1.00 | n.s. | 0.12 | 52.75 | 59.14 | -0.78 | n.s. |
| evm.TU.Scaffold_1_222469999.394 | 229 | 0.44 | 4.71 | 1.00 | n.s. | 0.34 | 48.89 | 108.25 | -0.07 | n.s. |
| evm.TU.Scaffold_1_222469999.1484 | 182 | 0.42 | 4.32 | 1.00 | n.s. | 0.30 | 87.99 | 72.76 | -1.02 | n.s. |
| evm.TU.Scaffold_1_222469999.4787 | 212 | 0.38 | 9.03 | 0.99 | n.s. | 0.18 | 63.17 | 69.29 | -1.15 | n.s. |
| evm.TU.Scaffold_3_155577901.2852 | 193 | 0.47 | 5.30 | 1.00 | n.s. | 0.21 | 56.40 | 62.97 | -0.80 | n.s. |
| evm.TU.Scaffold_1_222469999.5029 | 1048 | 0.41 | 5.12 | 1.00 | n.s. | 0.19 | 60.36 | 75.19 | -0.58 | n.s. |
| evm.TU.Scaffold_1_222469999.1143 | 207 | 0.43 | 4.22 | 1.00 | n.s. | 0.22 | 55.67 | 97.95 | -0.47 | n.s. |
| evm.TU.Scaffold_3_55577901.1386 | 217 | 0.34 | 6.12 | 0.99 | n.s. | 0.31 | 49.03 | 81.25 | -0.63 | n.s. |
| evm.TU.Scaffold_3_155577901.2294 | 369 | 0.46 | 5.06 | 1.00 | n.s. | 0.25 | 64.49 | 54.81 | -1.21 | n.s. |
| evm.TU.Scaffold_1_222469999.1464 | 461 | 0.41 | 6.05 | 1.00 | n.s. | 0.24 | 44.23 | 87.04 | -0.68 | n.s. |
| evm.TU.Scaffold_1_222469999.231 | 100 | 0.45 | 5.61 | 0.59 | n.s. | 0.36 | 46.90 | 107.37 | -0.35 | n.s. |
| evm.TU.Scaffold_1_222469999.29 | 245 | 0.43 | 4.68 | 1.00 | n.s. | 0.26 | 53.24 | 74.30 | -0.80 | n.s. |
| evm.TU.Scaffold_1_222469999.3878 | 331 | 0.45 | 7.53 | 1.00 | n.s. | 0.26 | 43.83 | 70.79 | -0.56 | n.s. |
| evm.TU.Scaffold_2_212802314.823 | 274 | 0.44 | 8.69 | 1.00 | n.s. | 0.18 | 55.28 | 64.98 | -1.07 | n.s. |
| evm.TU.Scaffold_3_55577901.923 | 217 | 0.43 | 4.40 | 1.00 | n.s. | 0.18 | 50.50 | 92.04 | -0.44 | n.s. |

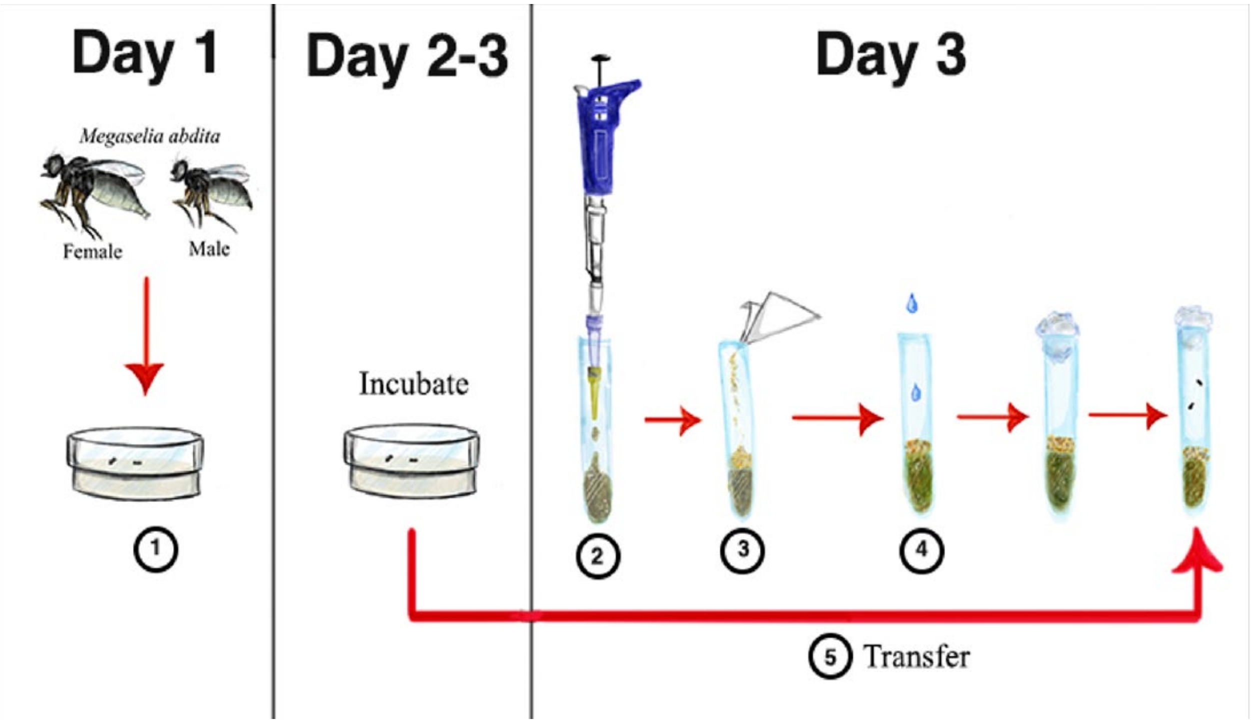


**Figure S11.** General overview of the protocol for the single crosses of *M. abdita*. Step 1: Virgin flies are collected and incubated in 2% agar-water gel plates for 2 days with a single male. Step 2: Food vials are prepared the day of transfer. Step 3: Once the agar has solidified, 0.1g additional food is added using weighing paper. Step 4: 200ul of water is added. Step 5: Flies are transferred into the food vials which are plugged with rayon and then incubated for ~21 days.

**Table S4**. All RNAseq sample names, accession numbers, metadata (from our study and NCBI).

| Sample Name | Developmental Stage | Source | BioSample | BioProject | SRA Run |
|---|---|---|---|---|---|
| M_abdita_F_RNA | Adult | Mahajan and Bachtrog. 2017. | SAMN06909123 | PRJNA385725 | SRR5559325 |
| M_abdita_M_RNA | Adult | Mahajan and Bachtrog. 2017. | SAMN06909124 | PRJNA385725 | SRR5559340 |
| RINSinlTBGRAAPEI-126 | Adult | Pauli et al. 2018. | SAMN03223122 | PRJNA267960 | SRR1695360 |
| mega-hiseq1 | Pooled Embryos (late blastoderm though early germband extension) | Jiménez-Guri et al. 2013. | SAMEA1572180 | PRJEB3172 | ERR196167 |
| Megaselia_454_ok | Pooled Embryos (late blastoderm though early germband extension) | Jiménez-Guri et al. 2013. | SAMEA1572181 | PRJEB3172 | ERR194165 |
| Mab_Stage_13 | Embryo - Stage 13 | This study | SAMN45895270 | PRJNA1200075 | SRR31763336 |
| Mab_Stage_15 | Embryo - Stage 15 | This study | SAMN45895271 | PRJNA1200075 | SRR31763335 |
| Mab_Stage_17 | Embryo - Stage 17 | This study | SAMN45895272 | PRJNA1200075 | SRR31763327 |
| Mab_Stage_1 | Embryo - Stage 1 | This study | SAMN45895273 | PRJNA1200075 | SRR31763321 |
| Mab_Stage_5 | Embryo - Stage 5 | This study | SAMN45895274 | PRJNA1200075 | SRR31763320 |
| Mab_Stage_8 | Embryo - Stage 8 | This study | SAMN45895275 | PRJNA1200075 | SRR31763318 |
| Mab_Stage_9 | Embryo - Stage 9 | This study | SAMN45895276 | PRJNA1200075 | SRR31763317 |
| Mab_Stage_10 | Embryo - Stage 10 | This study | SAMN45895277 | PRJNA1200075 | SRR31763316 |
| Mab_Stage_12 | Embryo - Stage 12 | This study | SAMN45895278 | PRJNA1200075 | SRR31763319 |
| Mab_L1 | Larva - 1st instar | This study | SAMN45895279 | PRJNA1200075 | SRR31763315 |
| Mab_L3 | Larva - 3rd instar | This study | SAMN45895280 | PRJNA1200075 | SRR31763334 |
| Mab_pupa | Pupa - day 1 | This study | SAMN45895281 | PRJNA1200075 | SRR31763333 |
| Dmel_Stage_13 | Embryo - Stage 13 | This study | SAMN45895282 | PRJNA1200075 | SRR31763332 |
| Dmel_Stage_15 | Embryo - Stage 15 | This study | SAMN45895283 | PRJNA1200075 | SRR31763331 |
| Dmel_Stage_16 | Embryo - Stage 16 | This study | SAMN45895284 | PRJNA1200075 | SRR31763330 |
| Dmel_Stage_17 | Embryo - Stage 17 | This study | SAMN45895285 | PRJNA1200075 | SRR31763329 |
| Dmel_Stage_1 | Embryo - Stage 1 | This study | SAMN45895286 | PRJNA1200075 | SRR31763328 |
| Dmel_Stage_5 | Embryo - Stage 5 | This study | SAMN45895287 | PRJNA1200075 | SRR31763326 |
| Dmel_Stage_8 | Embryo - Stage 8 | This study | SAMN45895288 | PRJNA1200075 | SRR31763325 |
| Dmel_Stage_9 | Embryo - Stage 9 | This study | SAMN45895289 | PRJNA1200075 | SRR31763324 |
| Dmel_Stage_10 | Embryo - Stage 10 | This study | SAMN45895290 | PRJNA1200075 | SRR31763323 |
| Dmel_Stage_12 | Embryo - Stage 12 | This study | SAMN45895291 | PRJNA1200075 | SRR31763322 |
